# Gene dosage effects of 22q11.2 copy number variants on in-vivo measures of white matter axonal density and dispersion

**DOI:** 10.1101/2025.05.29.656839

**Authors:** Rune Boen, Julio E. Villalón-Reina, Leila Kushan, Kathleen P. O’Hora, Hoki Fung, Nadine Parker, Ibrahim A. Akkouh, Dag Alnæs, Ruth O’Hara, Matthew John Marzelli, Lara Foland-Ross, Christina French Chick, Isabelle Cotto, Allan L. Reiss, Joachim Hallmayer, Paul M. Thompson, Ole A. Andreassen, Ida E. Sønderby, Carrie E. Bearden

## Abstract

22q11.2 deletion (22qDel) and duplication (22qDup) carriers have an increased risk of neurodevelopmental disorders and exhibit altered brain structure, including white matter microstructure. However, the underlying cellular architecture and age-related changes contributing to these white matter alterations remain poorly understood. Neurite orientation dispersion and density imaging (NODDI) was used on mixed cross-sectional and longitudinal data to examine group differences and age-related trajectories in measures of axonal density (i.e., intracellular volume fraction; ICVF), axonal orientation (orientation dispersion index; ODI) and free water diffusion (isotropic volume fraction; ISO) in 50 22qDel (n scans = 69, mean age = 20.7, age range = 7.4-51.1, 64.0% female) and 24 22qDup (n scans = 34, mean age = 21.6, age range = 8.3-49.4, 54.2% female) carriers, and 890 controls (n scans = 901, mean age = 21.9, age range = 7.8-51.1, 54.5% female). The results showed widespread gene dosage effects, with higher ICVF in 22qDel and lower ICVF in 22qDup compared to controls, and region-specific effects of the 22qDel and 22qDup on ODI and ISO measures. However, 22qDel and 22qDup carriers did not exhibit altered age-related trajectories relative to controls. Observed differences in ICVF suggest higher white matter axonal density in 22qDel and lower axonal density in 22qDup compared to controls. Conversely, differences in ODI are highly localized, indicating region-specific effects on axonal dispersion in white matter. We do not find evidence for altered developmental trajectories of axonal density or dispersion among 22q11.2 CNV carriers, suggesting stable disruptions to neurodevelopmental events before childhood.

## Introduction

22q11.2 copy number variants (CNVs) are rare, recurrent structural genetic variants that have profound impacts on neurodevelopment and confer increased risk for several developmental neuropsychiatric disorders, including autism spectrum disorder (ASD) and attention deficit/hyperactivity disorder (ADHD)(1–4). The 22q11.2 deletion (22qDel) is estimated to occur in one in ∼3700 live births(1), where 90% of cases are estimated to be due to de novo events(5). The 22qDel is also associated with a range of medical and neurobehavioral outcomes, including congenital heart defects, immune deficiencies, and intellectual disability(1,6,7), and confers a markedly increased risk of psychosis(1–4). The 22q11.2 duplication (22qDup) is estimated to occur in one in ∼1600 live births(1) and, conversely, confers a putative reduced risk of psychosis(4,8,9). Moreover, the 22qDup is more likely to be inherited rather than de novo, and may have a more variable clinical phenotype presentation(10,11) compared to the 22qDel. Despite their divergent risk for psychotic illness, both 22qDel and 22qDup are associated with elevated rates of ASD, ADHD, and intellectual disability compared to controls(1,3,12), and disrupt dosage-sensitive genes that play fundamental roles during neurodevelopment. Thus, in contrast to research on individuals with behaviorally defined psychiatric disorders, studying a molecularly confirmed CNV like the 22q11.2 locus provides a genetically traceable and hypothesis-driven probe of brain alterations in individuals with a high risk of developing brain disorders. Moreover, reciprocal CNVs (i.e., gain and loss of genomic material at the same locus) allows for investigation of converging and diverging neurodevelopmental pathways of brain alterations by examining gene dosage effects on brain phenotypes.

22q11.2 CNVs have been associated with medium to large differences in several measures of brain structure, including grey and white matter(13–16), suggesting altered cellular architecture. However, the complex neuronal composition and dispersion of the cortical mantle of the brain makes it challenging to study the cellular architecture in MRI-derived measures of grey matter, such as cortical thickness. In contrast, the white matter consists of predominantly axons, which can be probed using diffusion MRI (dMRI) in vivo. As water diffusion is restricted or hindered by biological tissue, dMRI can be used to generate measures of white matter microstructure that is influenced by the cellular architecture along white matter tracts(17–19). For instance, water molecules in an environment of dense and coherent white matter axons will diffuse along the primary direction of the axons, which has been commonly measured by diffusion tensor imaging (DTI)-derived metrics such as fractional anisotropy (FA, i.e., standardized value of anisotropic diffusion) and mean diffusivity (i.e., measure of diffusion in all directions).

A previous consortium-based DTI-study of 22qDel carriers reported an overall pattern of higher FA and lower diffusivity across most white matter tracts in 22qDel carriers compared to controls(13), which could reflect an increase in the cumulative cellular membrane circumference, increase in smaller tortuous axons(13), and/or higher density of axons with disproportionately small diameters(7, 14) compared to controls. Indeed, other studies have suggested other lines of evidence for increased axonal density in 22qDel carriers, such as excessive prenatal overgrowth of thalamocortical axons in a cellular model of the 22qDel(21) and a higher frequency of deep layer cortical neurons in a post-mortem examination of a 3-month-old infant with 22qDel(22). In contrast to 22qDel, only one prior small dMRI study has included 22qDup carriers(17). Despite the modest sample size, the results indicated widespread lower FA and higher diffusivity in 22qDup carriers compared to controls(23). Although the biological underpinnings of these differences are unknown, the opposing pattern in 22qDel and 22qDup carriers suggests a gene dosage effect, which may reflect divergent effects of the 22q11.2 CNV on the cellular architecture of white matter tracts. Indeed, the 22q11.2 locus is a hotspot for genes critical to white matter development, including axonal morphology and function in white matter such as regulation of axonal diameter, growth and branching (e.g., *ZDHHC8*(24,25)*, TBX1*(26)*, RTN4R/Nogo-66*(27)*, TXNRD2*(28)), as well as dendritic morphology and neuronal excitability (e.g., *DGCR8*(29,30)). In addition, both 22qDel and 22qDup affect expression of genes outside the 22q11.2 locus(21,29,31,32), including genes that influence axonal morphology(21). However, to our knowledge, no previous studies have directly compared axonal density in white matter tracts between 22qDel and 22qDup carriers to determine if a gene dosage effect exists. Moreover, such effects may be region-specific due to the unique developmental trajectories of white matter fibers. For example, limbic, commissural and projection fibers develop and mature earlier than association fibers(33–36). Indeed, it has been postulated that the nature of the white matter alterations in 22qDel carriers may differ between commissural and association fibers(13), potentially influenced by the maturational patterns of these fiber tracts. Specifically, the presence of commissural fibers are observed in the first trimester to second trimester(37–42), whereas the emergence of association fibers occurs later during the second trimester(38,39,42,43). The number of axons peak around birth for commissural fibers(44,45), followed by axonal elimination(45,46), axon growth and myelination from birth through adolescence (42,47,48). The association fibers are the least mature fibers during infancy(49) and are the last to mature, as indicated by developmental changes in DTI measures well into adulthood(36). Thus, it is plausible that the 22q11.2 CNV may have maturation-specific effects on commissural and association tracts.

Conventional DTI measures are inherently nonspecific to the underlying white matter cellular architecture(18,50,51) and can be influenced by alterations to axonal density and/or dispersion that can be driven by early (e.g., axonal proliferation or axonal pruning) or later (e.g., axonal diameter expansion) neurodevelopmental events. In contrast, neurite orientation dispersion and density imaging (NODDI) is a dMRI method that is better suited to test the hypothesis of altered axonal density in 22q11.2 CNV carriers, as it provides measures of the biophysical properties of white matter tracts, including axonal density (i.e., intracellular volume fraction, ICVF), axonal dispersion (i.e., orientation dispersion index, ODI), and free water diffusion/cerebrospinal fluid (i.e., isotropic water diffusion, ISO). It is important to note that NODDI generates metrics that provide indirect proxies of underlying microstructural properties and cannot directly disentangle the specific biological events (e.g., along-axon diameter variations and/or glial changes) that drive these signal differences(52). Nevertheless, as a novel technique that disentangles the composite signal of FA derived from DTI, NODDI metrics allow for more precise and biologically meaningful inferences about the underlying white matter architecture. Moreover, NODDI-derived measures have been found to correlate with their histological counterparts(53–55) and to be altered in several neuropsychiatric disorders, including psychosis and ASD(56). Thus, examining NODDI measures in 22q11.2 CNV carriers may be informative for detecting converging neurobiological patterns with idiopathic developmental neuropsychiatric disorders. In addition, ICVF has been found to be more sensitive to age-related differences in white matter microstructure compared to DTI-derived metrics during development(57). As such, this metric can yield novel insight into age-specific neurobiological events that underlie changes in white matter microstructure(58–60), such as prenatal proliferation of axons(44,45) and prolonged periods of axonal diameter expansion across childhood and adolescence(58). Thus, by mapping the neurodevelopmental trajectories of the axonal architecture across white matter regions, we can obtain important insight into the maturational mechanisms that underlie altered white matter observed in 22qDel and 22qDup carriers.

Here, we aim to characterize the effects of 22q11.2 CNVs on axonal density and axonal dispersion in white matter regions. We hypothesize gene-dosage effects on axonal density in white matter regions such that 22qDel carriers will exhibit higher and 22qDup carriers will exhibit lower axonal density, respectively. We also aim to investigate spatiotemporal effects of the 22q11.2 CNV on axonal architecture by examining age-related trajectories using mixed cross-sectional and longitudinal data, and by leveraging a large dataset of typically developing controls from the Human Connectome Project (HCP)(61,62). We reasoned that alterations to prenatal neurodevelopmental processes will manifest as robust group differences in NODDI measures, whereas alterations to later neurodevelopmental processes will be reflected by altered age-related trajectories in NODDI measures across childhood and adolescence.

## Methods

### Participants

A total of 964 participants with 1,004 scans were included in the analysis, including 50 22qDel carriers, 24 22qDup carriers and 890 age and sex matched typically developing controls (Table 1). The 22qDel carriers were recruited from either the University of California, Los Angeles (UCLA) or Stanford University, whereas the 22qDup carriers were recruited from UCLA only. The control group included unrelated healthy control participants recruited from UCLA and Stanford University, and healthy control participants derived from the HCP, including HCP-Development(61) and HCP-Aging(62) to establish robust developmental trajectories from childhood to adulthood (see sTable 1-2 for baseline characteristics across scanner sites, sTable 3 for binned age groups, sFigures 1-2).

**Table 1.**
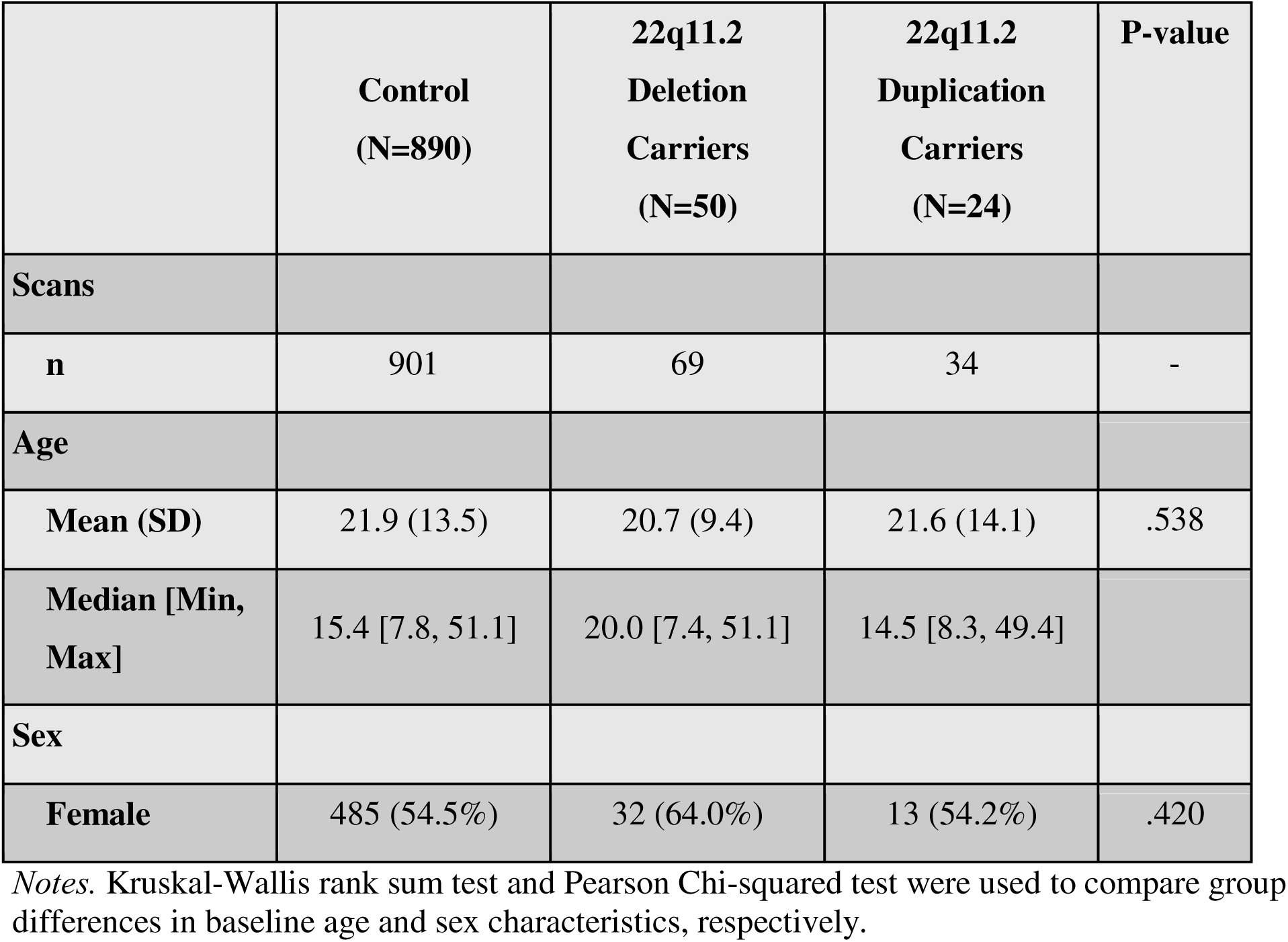
Participant baseline demographics.

|  | **Control**<br>**(N=890)** | **22q11.2**<br>**Deletion**<br>**Carriers**<br>**(N=50)** | **22q11.2**<br>**Duplication**<br>**Carriers**<br>**(N=24)** | **P-value** |
| --- | --- | --- | --- | --- |
| **Scans** |  |  |  |  |
| **n** | 901 | 69 | 34 | - |
| **Age** |  |  |  |  |
| **Mean (SD)** | 21.9 (13.5) | 20.7 (9.4) | 21.6 (14.1) | .538 |
| **Median [Min,**<br>**Max]** | 15.4 [7.8, 51.1] | 20.0 [7.4, 51.1] | 14.5 [8.3, 49.4] |  |
| **Sex** |  |  |  |  |
| **Female** | 485 (54.5%) | 32 (64.0%) | 13 (54.2%) | .420 |

*Notes.* Kruskal-Wallis rank sum test and Pearson Chi-squared test were used to compare group differences in baseline age and sex characteristics, respectively.

### Diffusion MRI acquisition, preprocessing and quality control

Multi-shell dMRI was acquired using a 3T Siemens Prisma scanner with the following parameters: TE = 89.2ms, TR = 3230ms, flip angle = 78°, slice thickness = 1.5mm, voxel size = 1.5mm isotropic obtained with 7 b = 0s/mm^2^ images, and 3 b = 200s/mm^2^, 6 b = 500s/mm^2^, 46 b = 1500s/mm^2^ and 46 b = 3000s/mm^2^ diffusion weighted volumes. The scans were acquired with both anterior to posterior and posterior to anterior phase encoding directions, resulting in a total of 216 volumes, following the HCP acquisition protocol(63). Manual inspections and statistical analyses were conducted to exclude participants with excessive head movement (see supplementary note 1 for details). We also included dMRI data derived from the HCP-D (https://www.humanconnectome.org/study/hcp-lifespan-development) and HCP-A (https://www.humanconnectome.org/study/hcp-lifespan-aging) as described elsewhere(61,62). All the raw dMRIs were denoised with LPCA and corrected for Gibbs ringing artifacts using DiPy (64,65); susceptibility induced distortions, head movement, and eddy currents using TOPUP and EDDY in FSL (66,67). Afterwards, we ran the diffusion preprocessing pipeline from HCP (https://github.com/Washington-University/HCPpipelines). NODDI was fitted using the dMIPY package (https://github.com/AthenaEPI/dmipy). The NODDI derived scalar maps were registered to the ENIGMA-DTI template(68) and average skeletonized ROI-based measures were calculated based on the Johns Hopkins University White Matter atlas(69) following the ENIGMA-DTI protocol (https://enigma.ini.usc.edu/protocols/dti-protocols/)(68), including anterior corona radiata (ACR), anterior limb of internal capsule (ALIC), body of corpus callosum (BCC), corpus callosum (CC), cingulum cingulate gyrus (CGC), cingulum hippocampus (CGH), corona radiata (CR), corticospinal tract (CST), extreme/external capsule (EC), fornix (FX), fornix (crus)/stria terminalis (FXST), genu of corpus callosum (GCC), internal capsule (IC), posterior corona radiata (PCR), posterior limb of the internal capsule (PLIC), posterior thalamic radiation (PTR), retrolenticular part of the internal capsule (RLIC), splenium of corpus callosum (SCC), superior corona radiata (SCR), superior fronto-occipital fasciculus (SFO), superior longitudinal fasciculus (SLF), sagittal stratum (SS), tapetum of the corpus callosum (TAP), uncinate fasciculus (UNC) (see sFigure 3 for a brain map). Due to repeated measures across scanner sites, we harmonized all of the NODDI measures using longCombat(70). Due to the focus on white matter regions, which mostly consist of myelinated axons(71), we define ICVF as a measure of axonal density, ODI as a measure of axonal dispersion and ISO as a measure of free water diffusivity(51).

### Statistical analyses

#### Group comparisons

We used generalized additive mixed models (GAMM) in R(72) to test for group differences in ICVF, ISO and ODI across 27 variables (24 regions of interest (ROIs) and 3 average measures). Here, we compared 22qDel carriers to controls, 22qDup carriers to controls, and 22qDel carriers to 22qDup carriers. In addition, we ran gene dosage analyses on NODDI measures by treating copy number as a continuous variable (22qDel = 1, control = 2, and 22qDup = 3). All group analyses were adjusted for the smoothed age effect using cubic regression splines, sex, and repeated measures through the inclusion of participant ID as random intercept. All GAMM analyses were fitted using default arguments, where k = -1 is set for the smoothed age terms and the use of restricted maximum likelihood when fitting all models. The standardized beta is reported and used as a measure of effect size. To account for multiple testing, we applied FDR correction across 75 (i.e., dependent variables) * 4 (i.e., three group analyses and one gene dosage analysis) = 300 comparisons. Finally, we conducted several sensitivity analyses to test the robustness of the results by including 1) participants from the same scanner site only, and by adjusting for 2) cerebral white matter volume, 3) intracranial volume or 4) cerebrospinal fluid volume (see supplementary note 2 for details).

#### Developmental trajectories

To examine the developmental trajectories of the NODDI measures, we examine the smoothed effect of age, with cubic regression splines using GAMM, in controls, 22qDel carriers and 22qDup carriers. All age-related analyses were conducted on site-harmonized data, and adjusted for the fixed effect of sex and the random effect of subject ID. First, we present the smoothed effect of age for the control group, along with the corresponding FDR-corrected p-values specific to the control group analysis, to establish typical developmental trajectories across white matter regions. The effective degrees of freedom (edf) represent the complexity of the smoothed age term, where values close to 1 indicate a linear relationship and higher values indicate non-linear relationships between chronological age and the response variable. We then compared the developmental trajectories of 22qDel and 22qDup carriers to the control group. A significant interaction term between the smoothed-age term and 22qDel or 22qDup indicates a difference in developmental trajectory compared to controls. To account for multiple testing, we applied FDR-correction across all the interaction terms (i.e., age*22qDel and age*22qDup).

## Results

### Gene dosage effects of the 22q11.2 CNV on intracellular volume fraction

Mean ICVF was significantly greater in 22qDel carriers compared to controls, and significantly greater across all regions except for FX and CGH. In contrast, 22qDup carriers showed significantly lower mean ICVF, and lower ICVF across all white matter regions compared to controls. Reflecting a similar pattern, there was a negative gene dosage effect of the 22q11.2 CNV (i.e., decreasing ICVF with increasing copy number) on mean ICVF and regional ICVF across all white matter regions (Figure 1, left panel, sTable 3). Sensitivity analyses revealed that scanner site, cerebral white matter volume, intracranial volume, and cerebrospinal fluid volume did not influence these results (sFigure 4, sTable 5-8).

**Figure 1.**
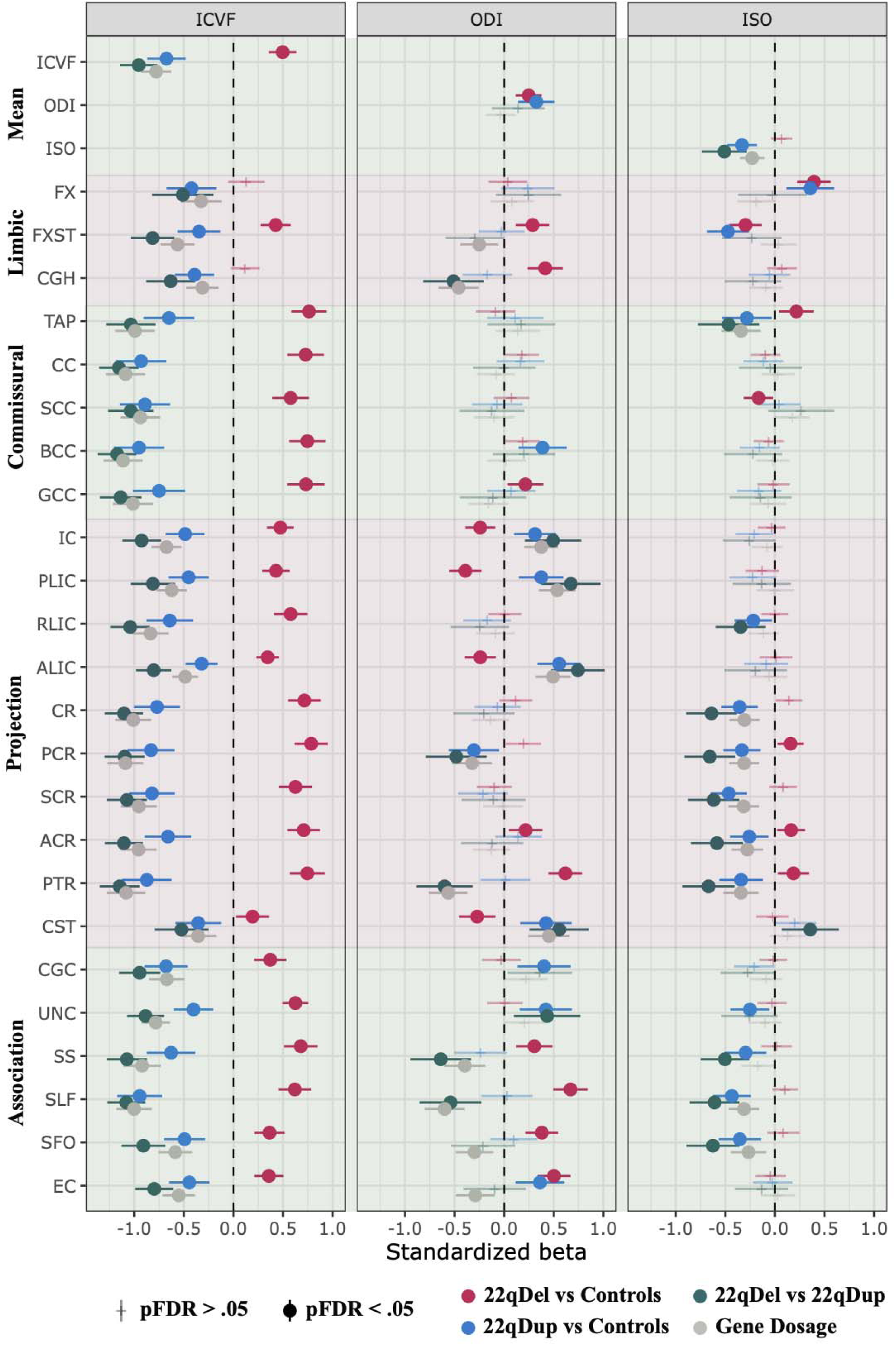
Point estimates of group differences in intracellular volume fraction (ICVF), orientation dispersion index (ODI) and isotropic volume fraction (ISO). The control group and the 22qDel group are used as reference groups for the group comparisons. Gene dosage effects are based on a continuous variable of copy number (i.e., 22qDel = 1, control = 2, and 22qDup = 3). E.g., Significant negative gene dosage effects indicate that an increased number of copies of genes in the 22q11.2 genomic locus is associated with lower values of the diffusion measures. ACR = anterior corona radiata, ALIC = anterior limb of internal capsule, BCC = body of corpus callosum, CC = corpus callosum, CGC = cingulum cingulate gyrus, CGH = cingulum hippocampus, CR = corona radiata, CST = corticospinal tract, EC = extreme/external capsule, FX = fornix. FXST = fornix (crus)/stria terminalis, GCC = genu of corpus callosum, IC = internal capsule, PCR = posterior corona radiata, PLIC = posterior limb of the internal capsule, PTR = posterior thalamic radiation, RLIC = retrolenticular part of the internal capsule, SCC = splenium of corpus callosum, SCR = superior corona radiata, SFO = superior fronto-occipital fasciculus, SLF = superior longitudinal fasciculus, SS = sagittal stratum, TAP = tapetum of the corpus callosum, UNC = uncinate fasciculus.

### Higher mean, but regionally variable effects, of 22q11.2 CNV on orientation dispersion index

Both 22qDel and 22qDup carriers showed greater mean ODI compared to controls; however, there were region-specific differences. 22qDel carriers had higher ODI in limbic (i.e., FXST, CGH), commissural (i.e., GCC) and association (i.e., SS, SLF, SFO, EC) fiber tracts, but also directionally variable effects on projection tracts (i.e., higher ODI in ACR and PTR; lower ODI in IC, PLIC, ALIC and CST) compared to controls. 22qDup carriers showed higher ODI in projection (i.e., IC, PLIC, ALIC, CST), commissural (i.e., BCC) and association (i.e., CGC, UNC, EC) fiber tracts, but also lower ODI in PCR, where projections fibers predominate, compared to controls (Figure 1, middle panel, sTable 3). The sensitivity analyses revealed the same pattern, albeit with fewer significant results; opposing effects of the 22qDel and 22qDup on ODI in projection (i.e., PTR, PLIC, IC) and association regions (i.e., SS and SLF) yielded the most robust effects (sFigure 5, sTable 5-8).

### Reduced free water diffusion in 22qDup carriers

There was no significant difference in mean ISO between 22qDel and controls. Regionally, however, 22qDel carriers exhibited higher ISO in a few projection tracts (i.e., PCR, ACR, PTR), and directionally variable effects on limbic (i.e., higher in FX; lower in FXST) commissural fibers (i.e., higher in TAP; lower in SCC) compared to controls (Figure 1, right panel). Sensitivity analyses revealed only higher ISO in the FX and PTR in 22qDel compared to controls after adjusting for intracranial volume, white matter volume (significant for FX and PTR), and cerebrospinal fluid volume (significant for FX; sFigure 3, sTable 3). 22qDup carriers showed lower mean ISO compared to controls, primarily driven by lower ISO in white matter regions predominated by projection fibers (RLIC, CR, PCR, SCR, ACR, PTR) and association fibers (UNC, SS, SLF, SFO, Figure 1, right panel). However, similar to the 22qDel carriers, 22qDup carriers also exhibited higher ISO in FX and lower ISO in FXST. Overall, sensitivity analyses yielded the same pattern of significant group-level differences between 22qDup and controls, except for the FX, UNC, RLIC and ACR that were no longer significant (sFigure 6, sTable 5-8).

### 22q11.2 CNV carriers show typical developmental trajectories in axonal density and dispersion

In controls, there were significant age effects on ICVF, ODI and ISO across all white matter regions from childhood to adulthood, except for ODI in the TAP (sTable 9). Most of the measures showed a non-linear age effect across white matter regions, except for ODI in ALIC, CST, EC, SFO, SLF and SS, which showed a linear age effect indicated by their effective degrees of freedom (i.e., edf = 1). The age-related trajectories of 22qDel and 22qDup carriers did not statistically differ from the age-related trajectories of the control group for the global and regional measures of ICVF (Figure 2), ODI (Figure 3) or ISO (Figure 4) after adjusting for multiple comparisons (sTable 10). Follow-up analyses also revealed that several group differences, most prominently in ICVF, are already evident in childhood and persist through adolescence and into adulthood (sFigure 7, sTable 11).

**Figure 2.**
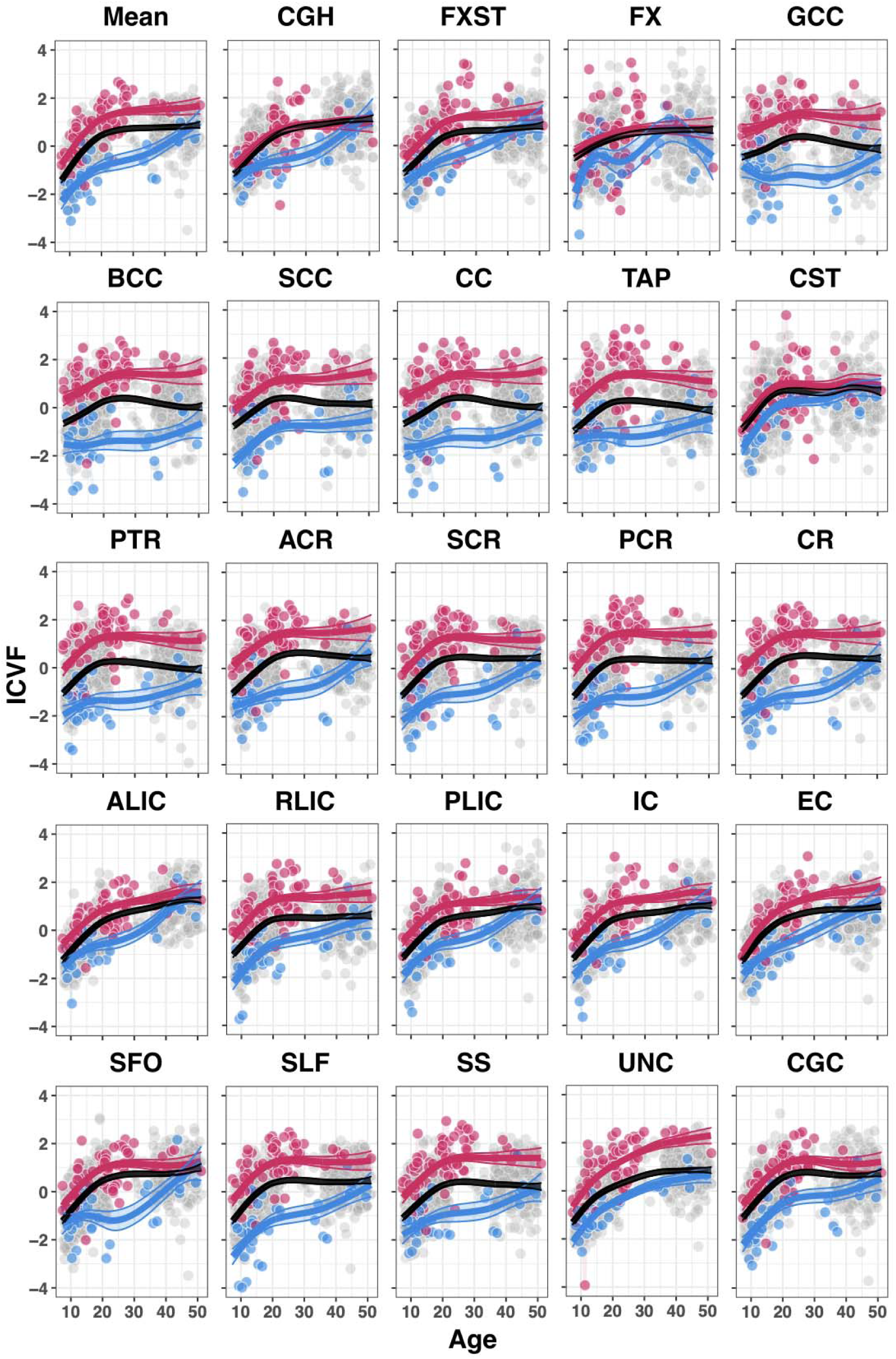
Developmental trajectories of intracellular volume fraction (ICVF) in global and regional white matter microstructure for 22q11.2 deletion carriers (red), controls (black) and 22q11.2 duplication carriers (blue). ACR = anterior corona radiata, ALIC = anterior limb of internal capsule, BCC = body of corpus callosum, CC = corpus callosum, CGC = cingulum cingulate gyrus, CGH = cingulum hippocampus, CR = corona radiata, CST = corticospinal tract, EC = extreme/external capsule, FX = fornix. FXST = fornix (crus)/stria terminalis, GCC = genu of corpus callosum, IC = internal capsule, PCR = posterior corona radiata, PLIC = posterior limb of the internal capsule, PTR = posterior thalamic radiation, RLIC = retrolenticular part of the internal capsule, SCC = splenium of corpus callosum, SCR = superior corona radiata, SFO = superior fronto-occipital fasciculus, SLF = superior longitudinal fasciculus, SS = sagittal stratum, TAP = tapetum of the corpus callosum, UNC = uncinate fasciculus.

**Figure 3.**
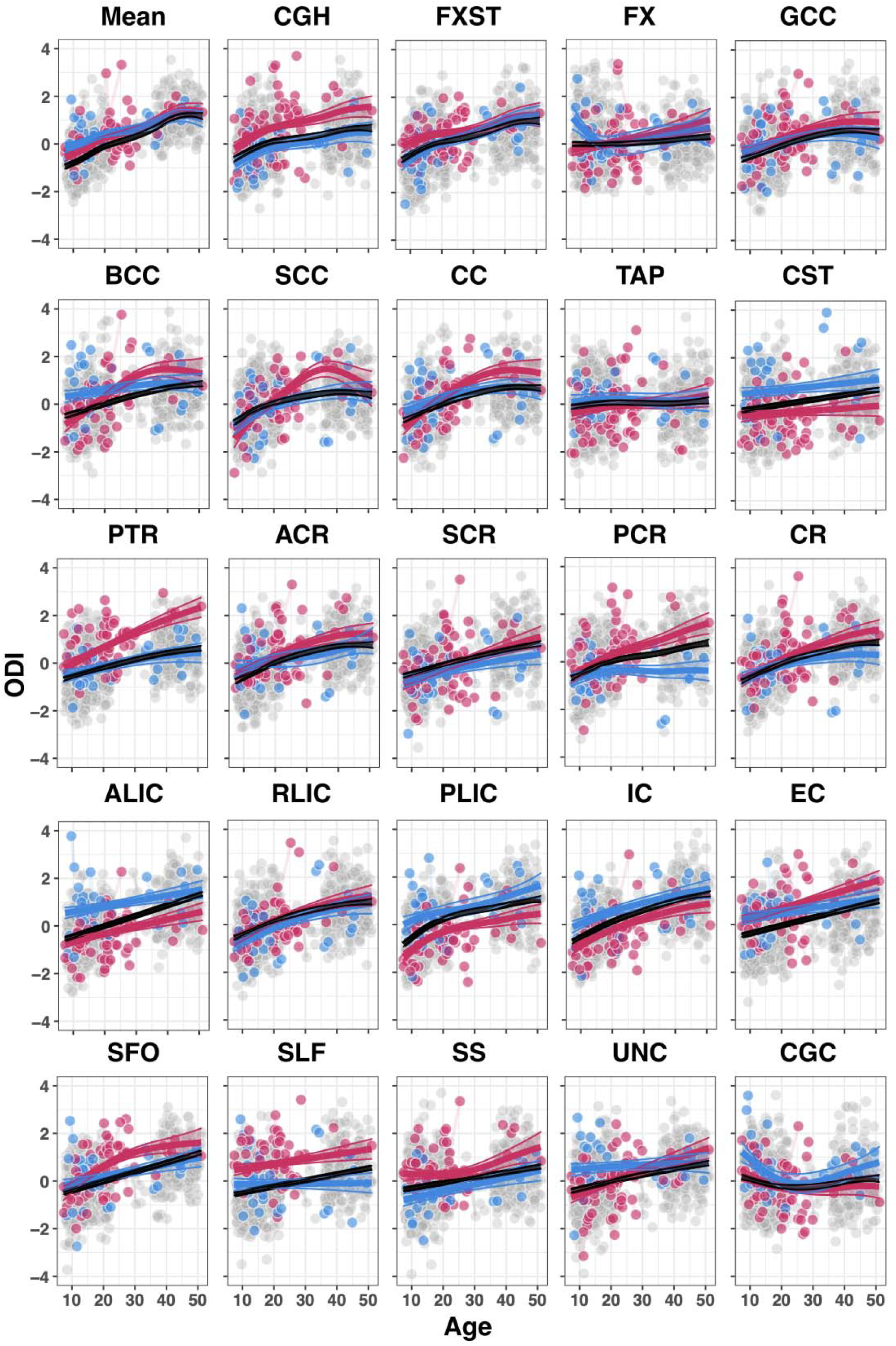
Developmental trajectories of orientation dispersion index (ODI) in global and regional white matter microstructure for 22q11.2 deletion carriers (red), controls (black) and 22q11.2 duplication carriers (blue). ACR = anterior corona radiata, ALIC = anterior limb of internal capsule, BCC = body of corpus callosum, CC = corpus callosum, CGC = cingulum cingulate gyrus, CGH = cingulum hippocampus, CR = corona radiata, CST = corticospinal tract, EC = extreme/external capsule, FX = fornix. FXST = fornix (crus)/stria terminalis, GCC = genu of corpus callosum, IC = internal capsule, PCR = posterior corona radiata, PLIC = posterior limb of the internal capsule, PTR = posterior thalamic radiation, RLIC = retrolenticular part of the internal capsule, SCC = splenium of corpus callosum, SCR = superior corona radiata, SFO = superior fronto-occipital fasciculus, SLF = superior longitudinal fasciculus, SS = sagittal stratum, TAP = tapetum of the corpus callosum, UNC = uncinate fasciculus.

**Figure 4.**
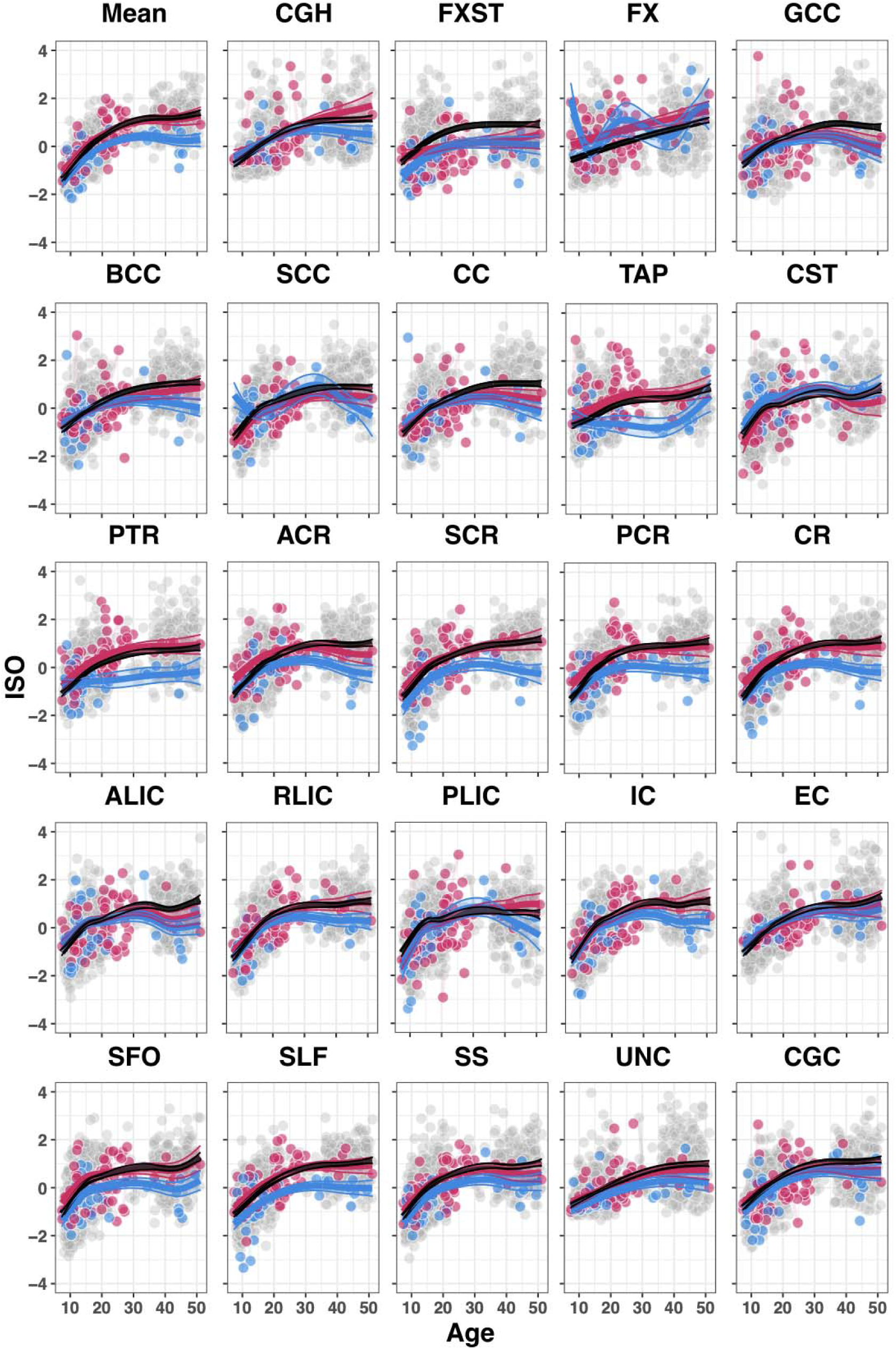
Developmental trajectories of isotropic volume fraction (ISO) in global and regional white matter microstructure for 22q11.2 deletion carriers (red), controls (black) and 22q11.2 duplication carriers (blue). ACR = anterior corona radiata, ALIC = anterior limb of internal capsule, BCC = body of corpus callosum, CC = corpus callosum, CGC = cingulum cingulate gyrus, CGH = cingulum hippocampus, CR = corona radiata, CST = corticospinal tract, EC = extreme/external capsule, FX = fornix. FXST = fornix (crus)/stria terminalis, GCC = genu of corpus callosum, IC = internal capsule, PCR = posterior corona radiata, PLIC = posterior limb of the internal capsule, PTR = posterior thalamic radiation, RLIC = retrolenticular part of the internal capsule, SCC = splenium of corpus callosum, SCR = superior corona radiata, SFO = superior fronto-occipital fasciculus, SLF = superior longitudinal fasciculus, SS = sagittal stratum, TAP = tapetum of the corpus callosum, UNC = uncinate fasciculus.

## Discussion

Hitherto, this is the largest study to examine biophysical dMRI-derived measures in brain white matter among 22q11.2 CNV carriers. The results showed widespread higher ICVF in 22qDel and lower ICVF in 22qDup compared to controls, where gene dosage at the 22q11.2 locus was associated with lower ICVF across all white matter regions, suggesting a strong effect of the 22q11.2 CNV on axonal density. Further, we found higher mean axonal dispersion (ODI for both 22qDel and 22qDup compared to controls, but also directionally variable effects of the 22q11.2 CNV on ODI across white matter regions; overall, having more copies of 22q11.2 genes was associated with lower ODI in regions where association, projection and limbic fibers predominate. However, we also observed that more copies at the 22q11.2 genomic locus were associated with higher ODI in IC and CST regions. In addition, we observed lower ISO in 22qDup carriers compared to controls in several white matter tracts where projection and association fibers predominate. Finally, we find that 22qDel and 22qDup carriers showed similar developmental trajectories to controls in all NODDI measures across childhood and adolescence.

The current study extends on the previous findings of higher FA in 22qDel(13) and lower FA in 22qDup(23) relative to typically developing controls, by focusing on measures of two of the main contributors to FA: axonal density and axonal dispersion(51). Across regions, we found robust evidence for higher axonal density in 22qDel carriers and lower axonal density in 22qDup carriers compared to controls, suggesting that axonal density is the main contributor to differences in FA alterations in white matter microstructure among 22q11.2 CNV carriers. It should be noted, however, that some white matter regions have previously been found to show lower (i.e., FXST, SLF, EC) or no difference in FA (i.e., RLIC, UNC, SS, PTR, SFO) in 22qDel carriers compared to controls(13). Here, we find that all these regions, except for the RLIC and UNC, were characterized by both higher axonal density and higher axonal dispersion in 22qDel carriers compared to controls. Thus, it seems that higher axonal dispersion counteracts the influence of higher ICVF, yielding lower or similar FA relative to controls in 22qDel carriers.

The greatest dosage effects of the 22q11.2 genomic region on ICVF were found in commissural tracts and the corona radiata, which consist of fibers in the corpus callosum and projection fibers from motor cortex, respectively. Such strong opposing effects on white matter architecture may be driven by mechanisms that are directly related to the deletion and duplication of genes within the 22q11.2 locus. However, the exact mechanisms underlying the dosage effect on these tracts are unknown, but it should be noted that the corpus callosum has been found to be larger in children with the 22qDel, which contrasts with the overall pattern of lower brain volume in 22qDel carriers(14,15). As corpus callosum is made up by subregions where the size of the axons typically differ in humans, e.g., the relative density of thick and thin axons (i.e., the axon diameter distribution) differs across the corpus callosum(73–75), it seems plausible that axonal overgrowth or axonal diameter expansion may underlie alterations to axonal density of commissural fibers and corpus callosum volume. Indeed, deletion of genes within the 22q11.2 locus (e.g., *ZDHHC8* and *RTN4R/Nogo-66)* has been linked to axonal growth and branching in mice models of the 22qDel(24,27). Others have shown that heterozygous deletion of the *TBX1* among mice can result in relatively more axons with small diameters and no large diameters in the fimbria compared to wild-type mice(26), which is in line with the predictions from previous diffusion MRI studies on 22q11.2 deletion carriers(13,20).

Our results also revealed higher ODI in 22qDel carriers for a few regions that are predominated by association fibers. Lower FA has previously been found in these white matter regions in 22qDel compared to controls(13). While lower FA can indicate both lower axonal density and/or higher axonal dispersion/crossing fibers(51), our result of increased ODI indicates that this is primarily driven by higher axonal dispersion in association tracts in 22qDel carriers relative to controls. Moreover, long association fibers typically originate from upper cortical layers and show protracted development compared to limbic and projection fibers(33–36), which may indicate that the white matter fibers originating from upper cortical layers show different morphology compared to fibers originating from deeper cortical layers. This interpretation aligns with several lines of previous research, including a mouse model of 22qDel showing fewer neural progenitors(76), shorter axons, and less dendritic branching in neurons from the upper cortical layers relative to wild type mice(28). The latter finding was observed after deletion of the *TXNRD2* gene, which also resulted in fewer long distance cortical connections and more local connections, which may yield higher axonal dispersion for long distance/association fibers in 22qDel carriers (e.g., lower alignment and/or rerouting through compensation of alternative or less direct paths). In addition, a post-mortem study reported lower frequency of neurons in the upper cortical layers in a 22qDel carrier(22), and a neuroimaging study reported altered axonal morphology (i.e., utilizing ultra-strong dMRI gradients sensitive to axonal morphology) and lower white matter volume in association tracts in 22qDel carriers compared to controls(20), supporting this interpretation.

Overall, the results point towards a divergent pattern of white matter microstructure between 22qDel and 22qDup carriers; however, there were a few points of convergence. Such diverging and converging effects are of particular interest in research on genomic structural variants, where gene dosage effects are often observed for neuroimaging derived features(16,23,77) but where the clinical phenotypic profile may converge (e.g., increased risk of ASD for both 22qDel and 22qDup carriers(12)). In the current study, we observed that a few regions exhibit similar directions of effect for both 22qDel and 22qDup, including higher ODI in EC, higher ISO in FX and lower ISO in FXST. These results indicate higher axonal dispersion in EC in both 22qDel and 22qDup, whereas the differences in FX and FXST may be driven by CSF partial volume in the voxels at the fornix and adjacent areas. White matter alterations in the external capsule have also been associated with idiopathic ASD (i.e., lower FA in twins with autism compared to control twins)(78) and symptom severity, and it is one of the white matter regions most susceptible to environmental factors in individuals with ASD(78). Moreover, a recent pilot study found within-subject white matter changes in the external capsule among toddlers with ASD after a behavioral intervention that improved verbal and communication skills(79), possibly implicating a role of the external capsule in individuals with or at risk for speech and communication problems. Our results suggest altered axonal circuitry in the external capsule may be implicated in the behavioral phenotypes of ASD commonly observed in 22q11.2 CNV carriers, although future research on the relationship between external capsule white matter structure and autistic phenotypes in 22q11.2 CNV carriers is required to understand this mechanism.

The 22qDel is also one of the greatest genetic risk factors for psychosis(4,80). A previous neuroimaging study has found evidence for lower axonal density in commissural and association fibers and higher axonal density and free water in the anterior thalamic radiation, predominated by projection fibers, among individuals with first episode psychosis(56). The differences in thalamic radiation converge with our results of higher ICVF and ISO in the PTR among 22qDel carriers compared to controls. A recent study reported thalamocortical axonal overgrowth in 22qDel thalamocortical organoids, likely due to elevated *FOXP2* expression as a trans-effect of the 22qDel(21). In addition, hyperconnectivity in thalamic circuitry has been reported among 22qDel carriers compared to controls(81–83), which has also been hypothesized as a potential biomarker for psychosis(84). It is also noteworthy that the 22qDup, carriers of a putative protective genetic factor for psychosis(4,8,9), showed the opposite pattern in our study, reflected by lower ICVF and ISO in the PTR. This may suggest that a higher axonal density of thalamic projection fibers, possibly due to thalamocortical axonal overgrowth, is implicated in the etiology of psychosis.

Finally, despite widespread group differences between 22q11.2 CNV carriers and controls, we did not find strong support for 22q11.2 CNV-dependent differences in the developmental trajectories of NODDI measures. Thus, the results do not support alterations in late maturational changes in axonal architecture, such as cumulative axonal diameter expansion. These findings contrast with the notably altered developmental pattern of resting-state functional connectivity observed in 22qDel carriers compared to controls(81,85). Concomitant with these findings, a parallel investigation in a mouse model of 22qDel found evidence for developmental changes in dendritic spine density(85). Taken together, these results suggest that changes in synapses and spine density may reshape brain circuits during adolescence, while altered axonal packing, as reflected by ICVF, may be established earlier. Still, it is important to note that several factors may contribute to these contrasting findings, including limited statistical power to detect interactions, high within-group variability, and/or imperfect biophysical specificity of the NODDI measures. Moreover, the observed alterations in axonal density may instead be reflective of atypical prenatal axonogenesis or lack of early axonal pruning in 22qDel and 22qDup carriers. This is in line with previous research on 22qDel-derived cortical spheroids exhibiting excessive prenatal overgrowth of thalamocortical axons(21) and neural progenitors that stay longer in the cell cycling state(86), which may be smaller in size, as indicated by reduced neurospheres derived from human induced pluripotent stem cells from 22qDel carriers(87). To speculate, the gene dosage effect of 22q11.2 genomic region on axonal density observed in our study may be reflective of a differential effect on gene expression levels of genes that are involved in axonogenesis, possibly through trans-regulatory mechanisms such as the mediating effect of *FOXP2* gene expression levels on 22qDel thalamic neurons(21). However, to our knowledge, no studies have utilized cellular models of the reciprocal 22qDup; such future studies are warranted to determine differential gene dosage effects on axonal development.

## Strengths and limitations

The current study includes a unique sample across a wide age-range consisting of both 22qDel and 22qDup carriers and a large typically developing control sample with multi-shell dMRI data, allowing for gene dosage and developmental analyses on metrics derived from advanced neuroimaging techniques. The use of advanced dMRI measures provides novel insight into the axonal architecture of the white matter microstructure in 22q11.2 CNV carriers, insight that cannot be achieved through conventional DTI measures(18,51). We also performed several sensitivity analyses demonstrating the robustness of the results of the group level analyses by analyzing individuals derived from the same scanner site only and adjusting for brain volumetric differences. The results revealed largely the same pattern, albeit with some group differences becoming attenuated due to reduction in sample size. It is also important to note that small sample sizes—due to the population prevalence of these CNVs—limit our ability to statistically detect small effects. Despite this, previous studies have found robust differences in white matter microstructure with smaller sample sizes than our study(20), which remains the largest study on biophysical white matter measures in 22q11.2 CNV carriers to date. Here, we also utilized GAMMs to capture nonlinear age effects, account for repeated measures and avoid biased group comparisons, especially important in instances where sample sizes are modest and age distributions differ between groups. Although we did not find evidence for altered developmental trajectories in 22q11.2 CNV carriers, we urge caution when interpreting these results, as the interaction effects may be more subtle in magnitude than the corresponding main effects and thus require greater power to detect. Thus, we cannot exclude the possibility of subtly altered developmental trajectories in NODDI measures among 22q11.2 CNV carriers compared to controls. However, because robust group differences in white matter microstructure between 22qDel carriers and controls are already evident in early adolescence(20), and neither our analyses nor a previous multi-site DTI study of 22qDel vs controls(13) show notable age-by-group interactions, there is currently no clear evidence that these group differences are driven by large maturational shifts during adolescence. Future studies with larger sample sizes, dense longitudinal data and methods that examine differences in smoothed confidence intervals(e.g., 15,86) may shed new light on possible age-dependent differences across the lifespan in 22q11.2 CNV carriers. Finally, despite the advantages of NODDI, it is still subject to the limitations inherent to dMRI, which is used to infer tissue microstructure by measuring water diffusion within a voxel of tissue. In addition, although NODDI measures have been shown to reflect their histological counterparts(53–55), it is important to note that biophysical models are emerging techniques that remain under active development(51,89–93). Our results also raise new questions. For instance, it is unclear if the altered axonal density in 22q11.2 CNV carriers is a consequence of an absolute difference in total neurons in the brain, altered axonal diameter distribution, or other mechanisms that alter space requirements in white matter tissue. Future studies utilizing both in-vitro cellular models and post-mortem examinations of 22q11.2 CNV carriers are warranted.

## Conclusions

The results of the current study provide new insights into the underlying neurobiology of the altered white matter microstructure in 22q11.2 CNV carriers, linking copy numbers at the 22q11.2 genomic locus to altered axonal density. However, we do not find evidence for altered age-related changes in axonal density or dispersion across childhood and adolescence, possibly reflecting atypical axonogenesis and/or axonal pruning during early neurodevelopment in 22q11.2 CNV carriers. Future studies are warranted to characterize the morphological features of white matter axons across fetal development, and to connect these developmental alterations to neuropsychiatric phenotypes.

## Acknowledgments

We thank Sergiu Pasca, M.D., for contributing to the study. This work used computational and storage services associated with the Hoffman2 Cluster which is operated by the UCLA Office of Advanced Research Computing’s Research Technology Group and servers operated by the USC Mark and Mary Stevens Neuroimaging and Informatics Institute (INI). Data from the UCLA and Stanford cohorts were supported by grants from the MCHRI Uytengsu-Hamilton 22q11 Neuropsychiatry Research Award (UH22QEXTFY22-04) and NIH 1R21MH116473-01A1, R37MH085953, 1U01MH119736-01 (to CEB), R01MH100900 (to JH), and R01MH129858 (to CEB, PMT, IES, OAA). Research reported in this publication was supported by the National Institute Of Mental Health of the National Institutes of Health under Award Number U01MH109589/ U01AG052564 and by funds provided by the McDonnell Center for Systems Neuroscience at Washington University in St. Louis. The HCP-Development 2.0 Release data used in this report came from DOI: 10.15154/1520708. The HCP-Aging 2.0 Release data used in this report came from DOI: 10.15154/1520707.

## Conflict of Interest

OAA is a consultant for Cortechs.ai and Precision Health and has received speaker’s honoraria from Lundbeck, Janssen, Otsuka, Lilly and Sunovion. The remaining authors have no conflicts of interest to declare.

Supplementary information is available at MP’s website.

